# Revealing the Prevalence of Suboptimal Cells and Organs in Reference Cell Atlases: An Imperative for Enhanced Quality Control

**DOI:** 10.1101/2024.04.18.590104

**Authors:** Tomàs Montserrat-Ayuso, Anna Esteve-Codina

## Abstract

The advent of droplet-based single-cell RNA-sequencing (scRNA-seq) has dramatically increased data throughput, enabling the release of a diverse array of tissue cell atlases to the public. However, we will show that prominent initiatives such as the Human Cell Atlas, Tabula Muris, and Tabula Sapiens contain a significant amount of contamination products (frequently affecting the whole organ) in their data portals due to suboptimal quality filtering. Our work addresses a critical gap by advocating for more stringent quality filtering, highlighting the imperative for a shift from existing standards, which currently lean towards greater permissiveness. We will show the importance of incorporating cell intronic fraction in quality control -or MALAT1 expression otherwise- showcasing its informative nature and potential to elevate cell atlas data reliability. In summary, here, we unveil the hidden intronic landscape of every tissue and highlight the importance of more rigorous single-cell RNA-sequencing quality assessment in cell atlases to enhance their applicability in diverse downstream analyses.

## Introduction

With the rapid advancement of single-cell RNA-seq technologies and the launch of global collaborative initiatives that contributed to the release of the human and mouse reference cell atlases, there is an unprecedented surge in data production. The Tabula Sapiens atlas, funded by the Chan Zuckerberg Initiative and published in Science^1^, comprises 500,000 human cells from 24 organs and the Tabula Muris atlas, published in Nature^2^, is a compendium of single cell transcriptome data from the model organism Mus musculus, containing nearly 100,000 cells from 20 organs and tissues. The Human Cell Atlas (HCA) Data Portal (https://data.humancellatlas.org/) houses data from over 50 million cells across 200 publications. These atlases represent a new resource to molecularly define cell types, organs, and tissues, and serve as an abundant asset to the single cell research community. However, a significant challenge has surfaced within these data repositories, as they have yet to attain the status of high-quality human and mouse reference cell atlases. We reveal that a significant number of the available organs include a substantial number (up to 85 %) of low-quality cells. This issue underscores a long-standing concern within the single-cell community that has, until now, not received the attention it deserves, which is the difficulty of obtaining high-quality scRNA-seq samples from certain solid tissues. The sample storage conditions, and tissue dissociation protocols usually impact cell viability resulting in varying degrees of contamination coming from damaged and dying cells.

Until we do not reach a solution for improved sample dissociation or accurate enrichment of intact cells prior to sequencing, the current strategy is to bioinformatically filter out low-quality cells before conducting any subsequent analyses. Widely adopted data processing workflows for droplet-based single-cell technologies, such as the Cell Ranger software suite^3^, distinguish between cells and empty droplets based on total UMI counts. After this step, Cell Ranger uses a variation of the EmptyDrops algorithm^4^, which uses the RNA profile of each remaining barcode to further separate cells from background noise. Additional filtering of sub-optimal cells is further implemented based on best practices analysis protocols and tutorials^5,6^ once the matrix of gene counts per cell is obtained, removing cells with low number of UMIs and high mitochondrial (MT) content (apoptotic cells). Accidentally, more than one cell ends up encapsulated in a drop. Doublets or multiplets should exhibit higher mRNA content than individual cells and sometimes present incompatible cell type markers; these can be removed with specific programs for doublet detection^7,8^. This procedure for eliminating low-quality cells may be effective in tissues easy to dissociate or cell lines where cells do not undergo aggressive treatment. However, it proves insufficient for obtaining high-quality cells from challenging tissues, such as heart, liver, or kidney, where the treatment to dissociate tissue cells is highly aggressive, damaging most of the cells. Ambient RNA originating from highly abundant lysed cells can be also inadvertently encapsulated in cell-free droplets or swarm around in cell-containing droplets. Several computational tools have been developed to mitigate the impact of this ambient RNA^9–11^. Others^12^, propose incorporating the nuclear fraction (intronic content) as a quality metric to identify and differentiate intact cells from cellular remnants (cytosolic or nuclei debris). Apart from very specific cell types such as erythrocytes and platelets, which lack a nucleus, all cells, once their mRNA is sequenced, should have a significant fraction of reads mapped to introns. Intron-absent cells likely represent empty droplets or cytosolic debris. Similarly, cells with an excessively high proportion of intronic reads suggest lysed cells with nuclei but some depletion of the cytosol. Despite these advancements, the practices of ambient RNA removal and intronic content calculation remain underutilized and relying solely on one of these methods is insufficient.

Often, these filters are applied uniformly to all cells in the sample, the biological or phenotypic variability among cell types is not considered. The mRNA content of a cell can vary for various reasons, such as cell size, cell cycle state, or cell type. In a complex tissue like the liver, a range of cell types exists, from binucleated hepatocytes to less complex cells like neutrophils (usually with low nUMIs and accidentally removed from the analysis). Similarly, metabolically active cells such as hepatocytes or cardiac cells may have a high content of mitochondrial transcripts without necessarily indicating that these cells are undergoing apoptosis. This can lead to two scenarios: either the filters are very strict, causing less complex cells to be eliminated along with poor-quality cells, or the filters are too lenient, allowing many poor-quality cells to pass through and be treated as genuine tissue cells. Most manuals and expert recommendations advise against strict quality control, suggesting instead to review the filtered cells after annotation^6,13^. Probably for this reason, as we will demonstrate in this study, many analyses of published scRNA-seq datasets include low-quality cells inappropriately. This issue is particularly serious because these cells often cluster closely with intact cells and are assigned to a specific cell type, or more critically, to a supposed new cell subtype. Therefore, new procedures are needed to identify and remove these cells from the analysis to avoid misleading conclusions.

In this study, we thoroughly reviewed the cell quality of publicly available datasets in reference atlases detecting a non-negligible number of low-quality cells often mistakenly designated as a distinct cell subtype with unique characteristics. We also show that sometimes the full organ is totally damaged. Empty droplets and cell debris could have been easily detected and removed from the analysis directly using the intron content metric. Paramountly, our study challenges the conventional association of MALAT1 with poor-quality datasets, clarifying its role as a nuclear marker constitutively expressed in all nucleated cells and considering it as a relevant quality metric.

Here, we completely reanalyzed four datasets using the raw data (from fastq) whose publications have had a significant impact on the single-cell community: a kidney dataset (10x Genomics and Smart-seq2), which has received (at the time of writing this manuscript) more than 1400 citations and is part of the Tabula Muris consortium^2^; a liver dataset, which has received more than 780 citations and is part of the Human Cell Atlas^14^; and a retina dataset, which has received more than 190 citations and is also part of the Human Cell Atlas^15^. Otherwise, the processed data was used to explore a mouse kidney cell atlas recently published by Novella-Rausell et al.^16^, the Tabula Muris Senis dataset and the Tabula Sapiens dataset, with more than 140000, 356000 and 480000 cells respectively, confirming the value of MALAT1 expression in the absence of the intron fraction quality metric.

In summary, our study emphasises the necessity of exceeding conventional quality metrics prevalent in many public single-cell RNA-seq studies to achieve elevated standards of quality. This is crucial in ensuring the dependability of cell atlases, thereby facilitating their effective utilisation in diverse downstream analyses.

## Materials and methods

### Collecting raw data from different sources

The raw data for the kidney dataset from Tabula Muris (10x Genomics and Smart-seq2 subsets) were downloaded from SRA with accession numbers SRX3791768, SRX3791769, SRX3791776 and SRX3607047. For the liver dataset, the raw data were downloaded as bam files from SRA with accession numbers SRR7276474, SRR7276475, SRR7276476, SRR7276478 and SRR7276477. Raw data as bam files for the retina dataset were downloaded from the Human Cell Atlas data portal in the following link https://explore.data.humancellatlas.org/projects/-8185730f-4113-40d3-9cc3-92927178 <u>4c2b</u>. The bam files of the kidney and retina datasets were converted to fastq files using the *bamtofastq* tool from the 10x Genomics Cell Ranger (v7.0.1) pipeline.

### Processing of sequencing data

For the 10X Genomics data, the *count* tool from Cell Ranger was used to generate the gene expression matrix for each cell for every fastq file. The unfiltered count matrices for every dataset were loaded into R (v4.2.0) and only the barcodes used by the original authors were kept (see below). Then we merged the different matrices of the same dataset to generate a single Seurat^17^ (v4.3.0) object for each dataset. The reads from Smart-seq2 technique were aligned to the gencode.vM19 mouse genome using STAR^18^ (v2.7.9a). Gene counts were produced using HTSeq^19^ (v2.0.2) with parameters “-t exon”, “-s no”, “-m intersersection-nonempty”,”-f bam” and “-i gene_name”. The introns content for each cell was obtained by parsing the output from Qualimap^20^ (v2.2.1) using a custom Python script. Finally, count matrices for each cell were merged into a single R data.frame and a Seurat object was created.

The expression values of each dataset were normalized with standard library size scaling and log transformation using Seurat’s NormalizeData() function with default parameters. The 3000 most variable genes were detected using the variance-stabilizing transformation selection method in Seurat (FindVariableFeatures() function). Then, we scaled the most variable genes using the ScaleData() function from Seurat with default parameters. From the standardized data, we calculated the principal components using the Seurat’s function RunPCA() and used the same number of principal components (pc) as the original authors used if reported or based on the respective elbow plots to generate the UMAP coordinates for further visualization of the dataset. Specifically, we used the first 40 pcs in the 10X Genomics kidney dataset from Tabula Muris, the first 30 pcs in the liver dataset, the first 20 pcs in the retina dataset and the first 30 pcs in the Smart-seq2 kidney dataset.

The original cell annotations, as well as the barcodes used by the authors were taken from different places depending on the dataset. For the Tabula Muris datasets, the Seurat object with the barcodes and annotations was downloaded following its web site instruction (https://tabula-muris.ds.czbiohub.org/). The annotations and barcodes used by the liver dataset’s authors were downloaded from GEO accession number GSE115469. Finally, the authors of the retina dataset kindly shared with us the cell metadata with the barcodes and cell annotations they used in their analysis.

### Collecting Seurat objects from CELLxGENE

The Seurat R object of the kidney cell atlas dataset from Novella-Rausell et al. (2023), as well as the whole Tabula Muris Senis, Tabula Sapiens and the Tabula Sapiens dataset for the kidney, heart and prostate subsets, as well as for the whole Tabula Sapiens dataset were downloaded from CELLxGENE: https://cellxgene.cziscience.com/collections/92fde064-2fb4-41f8-b85c-c6904000b859 for the Novella-Rausella et al. (2023) kidney cell atlas dataset, https://cellxgene.cziscience.com/ <u>collections/0b9d8a04-bb9d-44da-aa27-705bb65b54eb</u>for the Tabula Muris Senis dataset and https://cellxgene.cziscience.com/collections/e5f58829-1a66-40b5-a624-9046778e74f5 for the Tabula Sapiens datasets. We would have liked to be able to access the Tabula Sapiens raw data, which are supposed to be available upon request, but we never got an answer from the authors.

### Calculation of the intronic content and MALAT1 metrics

The introns content for each cell was determined using the function nuclear_fraction_tags() from the DropletQC R package. We used the function identify_empty_drops() from DropletQC with default parameters, which automatically determine those cells with extremely low introns content or moderately low introns content along with reduced number of reads/UMIs, classifying each barcode as “empty droplet” or “cell”. While exploring the MALAT1 expression in the Tabula Sapiens dataset, we classify the cells as “MALAT1-” if they did not have any MALAT1 count, and “MALAT1+” otherwise.

### Comparison of the mitochondrial percentage between single-cell and single-nucleus RNA-seq

For the Novella-Rausell kidney dataset, we divided the single-cell and single-nucleus barcodes, and, for each technology, we studied the distribution of the mitochondrial read percentages depending on the amount of MALAT1 expression in each cell. The “cytosolic debris” was formed by those cells with exactly zero MALAT1 normalized counts; the “cells” group was made up by cells with more than 0 and less or equal to 3.5 MALAT1 normalized counts; and the “nuclei enriched” group corresponded to barcodes with more than 3.5 MALAT1 normalized counts.

## Results

### A significant proportion of cells with null intronic fraction in reference atlases

The first scRNA-seq dataset inspected belongs to the Tabula Muris project, specifically the dataset obtained from microfluidics (10x Genomics) of the kidney. The original data already processed by the authors can be interactively explored on the Tabula Muris data portal: https://tabula-muris.ds.czbiohub.org/. In the original study, they applied the standard quality control, cells with fewer than 500 genes (nFeatures) or fewer than 1000 captured UMIs (nUMIs) were discarded. In total, 2781 cells passed the filters and were deemed suitable for downstream analysis, where they were classified into 8 different cell types. In Figure 1A, it can be observed that among the cells used by the authors for the analyses, 5 clusters with virtually zero nuclear fraction appear. Significantly, the epithelial cells of the proximal straight tubule are arranged in two nearly specular clusters, one with high and the other with null intron content. Clusters with no intronic reads were identified as containing empty droplets by DropletQC. At least, a total of 835 (precisely 30.03% of the cells in the dataset) cells from 4 different cell types (capillary endothelial cells, collecting duct epithelial cells, loop of Henle ascending limb epithelial cells and proximal straight tubule epithelial cells) should have been discarded or flagged during the initial quality control (Table 1).

**Figure 1:**
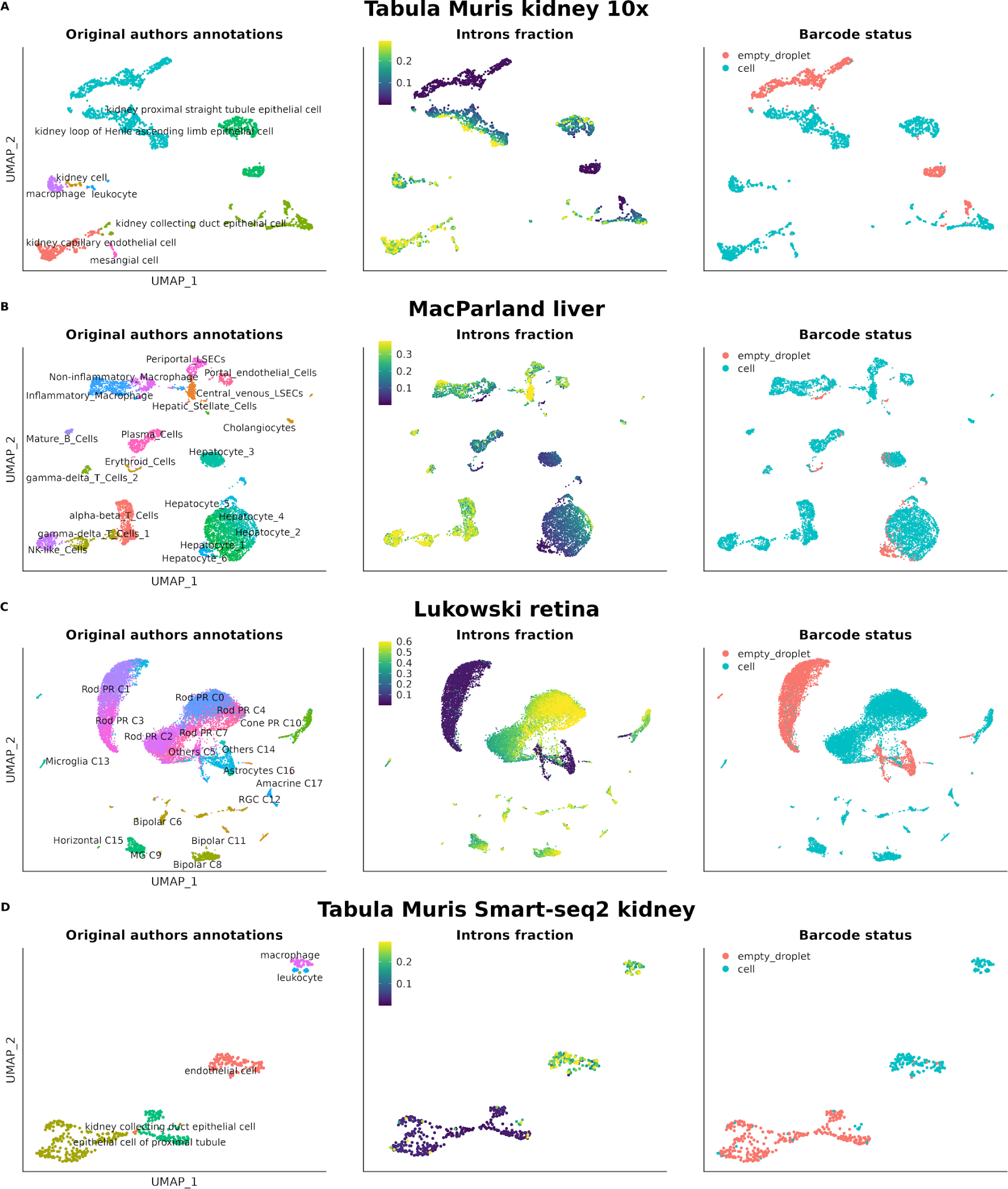
UMAPs showing cell type annotation, nuclear fraction and empty droplet/cell classification. A) Kidney 10X Genomics, B) Liver 10X Genomics, C) Retina 10X Genomics, D) Kidney Smart-Seq2.

**Table 1:**
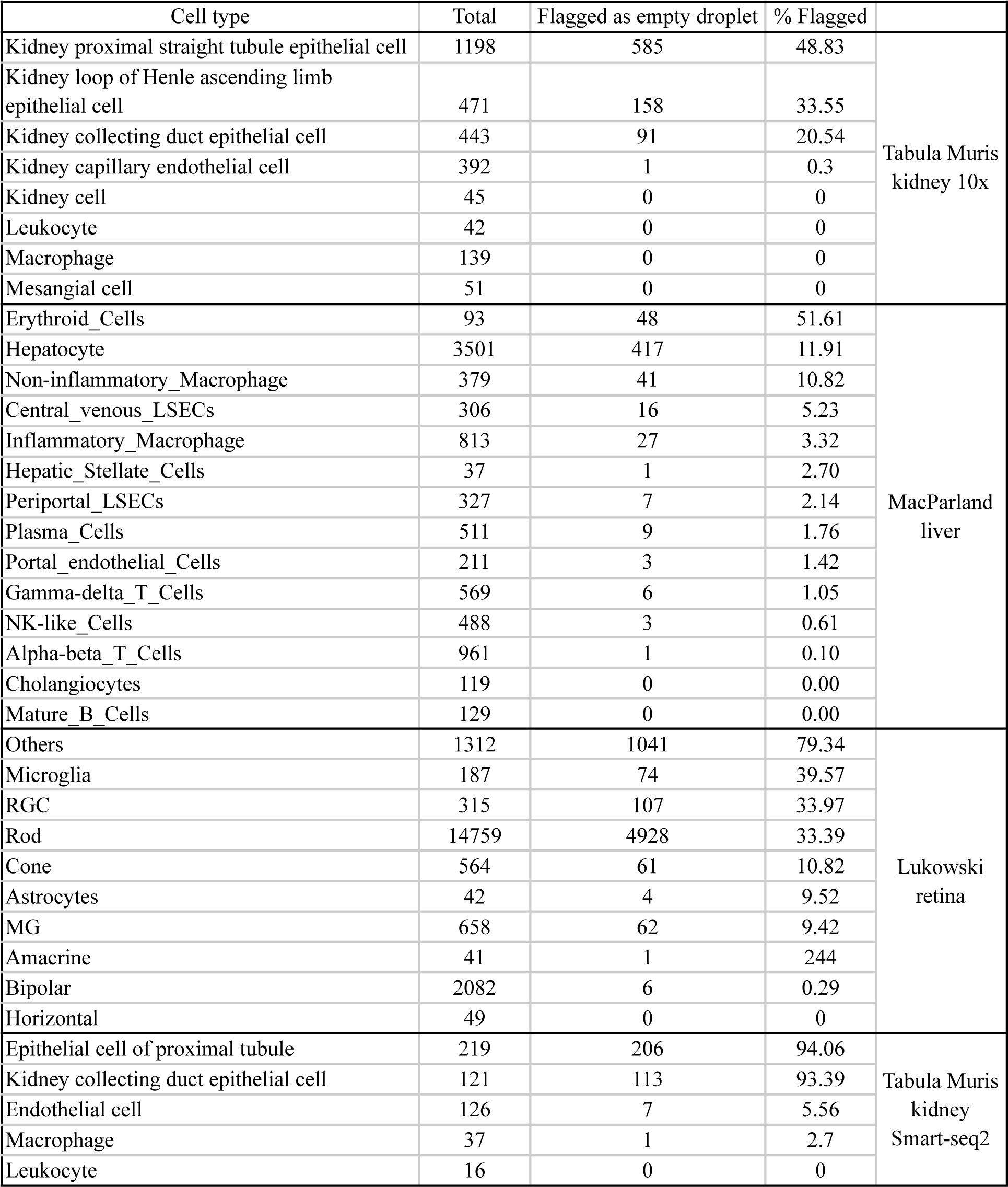
Empty droplet flag percentage per cell type.

We also inspected the data from the first cell atlas of the human liver, which was published by MacParland et al.. The quality control conducted in this study, once again, was the standard; cells with more than 1500 UMIs and less than 50% of UMIs mapped to mitochondrial genes passed the initial filtering. In total, 8444 cells, classified into 9 cell types and 20 different clusters, form this atlas. In this dataset, the majority of hepatocytes exhibit very low intronic content with their cluster showing a gradient in this regard. Interesting to note, the cluster annotated by the authors as “Hepatocyte_6” consists of cells without intronic content and, after running DropletQC, they were identified as empty droplets. On the contrary, non-parenchymal cells show much higher contents. At least (if not all), 417 hepatocytes, constituting 11.91% of all hepatocytes, should not have passed the quality control and be removed or flagged (Table 1). Other small clusters (from endothelial cells, macrophages and erythroids) were also identified as empty droplets by the extremely reduced amount of nuclear fraction (Table 1). Erythrocytes, as expected for anucleated cells, also appear as cells with a residual amount of intronic content (Figure 1B).

Finally, we analyzed the single-cell atlas of the adult human retina published by Lukowski et al. in 2019. This atlas can be accessed on CELLxGENE by following this link: https://cellxgene.cziscience.com/e/d5c67a4e-a8d9-456d-a273-fa01adb1b308.cxg/. The authors of this study performed traditional quality control, classifying cells with fewer than 200 or more than 2500 detected genes and expressing more than 10% of mitochondrial genes as low-quality cells. A total of 20,009 cells passed these filters and constitute this atlas. In the publication, a total of 10 different cell types were reported across 18 clusters based on the transcriptional state of the cells. As observed in Figure 1C, there are 4 clusters formed by droplets without immature mRNA, classified as empty droplets after running DropletQC. Interestingly enough, rod cells appear in two big clusters with high (C0, C2, C4, C7) and low intron content (C1, C3), and the cells classified as “Others” also have null intron content. Thus, at least 6284 cells from 9 different cell types, representing 31.41 % of the cells in the dataset, should not have been included in the analyses (Table 1).

The presence of cells with null intron content was also observed in the Smart-seq2 dataset of the Tabula Muris kidney, and also were flagged as “empty droplet” by DropletQC. Here, similar quality control as for the 10x Genomics data was applied by the original authors: cells with fewer than 500 genes or 50000 reads were discarded. The most affected cells coincided with those from the same tissue but obtained with the droplet-based technology from 10x Genomics (Figure 1D).

### MALAT1 can be used as a surrogate for cellular nuclear content

The cells without immature mRNA and annotated as empty droplets by DropletQC in the kidney, liver, and retina datasets, also exhibited very low or no expression of the long non-coding RNA MALAT1 (Figure 2A-D). All clusters with a null content of immature mRNA (except for cells annotated as erythrocytes, which naturally should not contain introns) displayed this characteristic. The correlation of MALAT1 expression and introns content is quite high, yielding Pearson correlations of 0.83 for Tabula Muris kidney 10x Genomics; 0.78 for MacParland liver; 0.92 for Lukowski retina; and 0.91 for Tabula Muris kidney Smart-seq2 (all p-values < 2.2×10^-16^).

**Figure 2:**
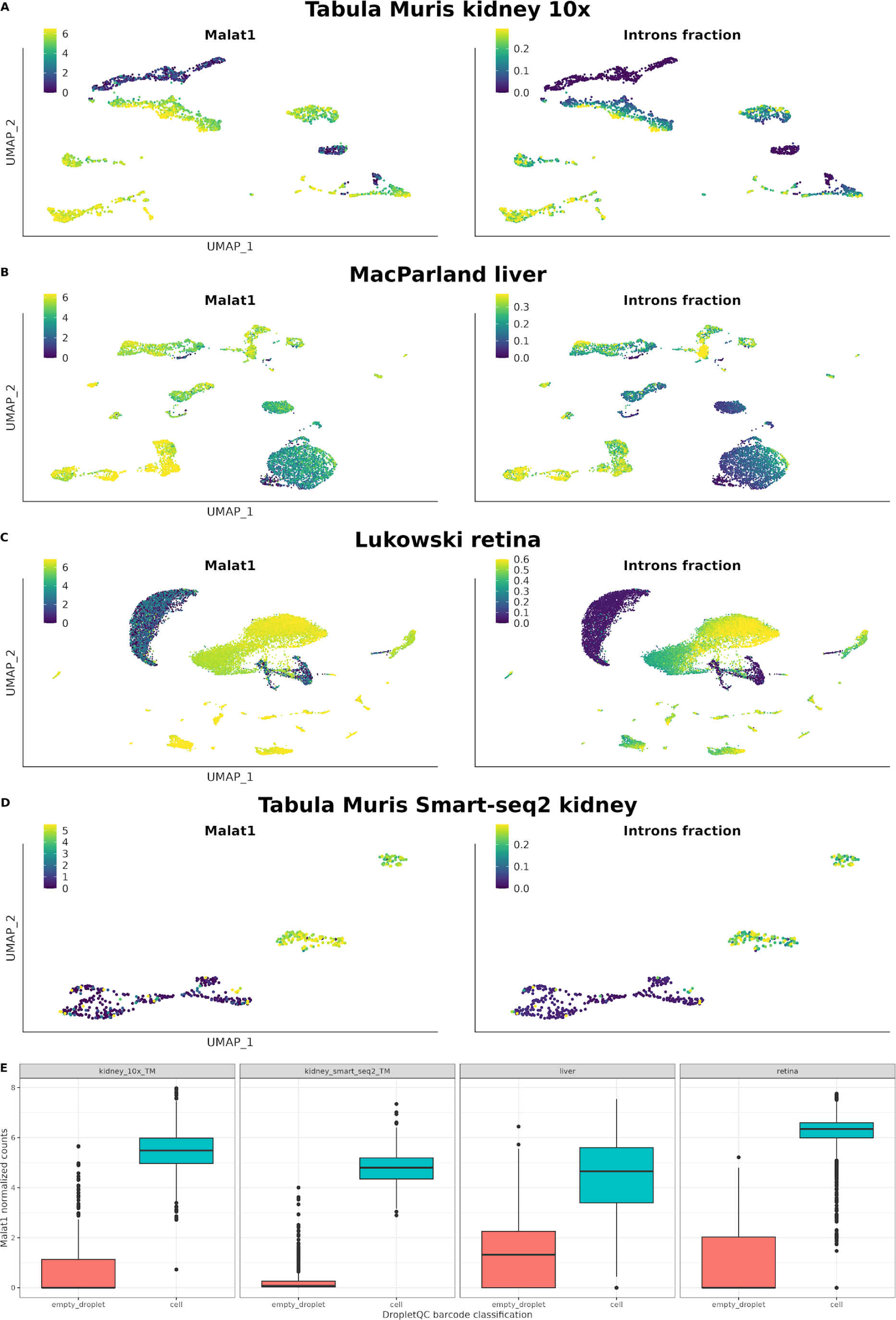
A-D) UMAPs showing the relationship between intron content and MALAT1 expression. E) Boxplots of MALAT1 expression in empty droplets and cell subsets.

Using the MALAT1 expression as a proxy of the intron fraction, we explored a recent publicly available kidney atlas dataset from Novella-Rausell et al. (2023), where we only got access to the count matrix and we could not calculate the percentage of intronic reads per cell. In this study, the authors constructed a cell atlas from 59 public kidney datasets from 8 different studies. After filtering cells based on the number of UMIs and the percentage of reads mapped to the mitochondrial genome, a total of 141,401 cells were considered suitable for inclusion in the atlas and for conducting relevant analyses. This dataset can be interactively explored on CELLxGENE through this link: https://cellxgene.cziscience.com/e/42bb7f78-cef8-4b0d-9bba-50037d64d8c1.cxg/. It consists of samples obtained from both single-cell RNA-seq and single-nucleus RNA-seq techniques. Single-cell samples showed many MALAT1-cell clusters, as opposed to single-nucleus samples (Figure 3A-C). This is consistent with the fact that MALAT1 is confined in the nucleus.

**Figure 3:**
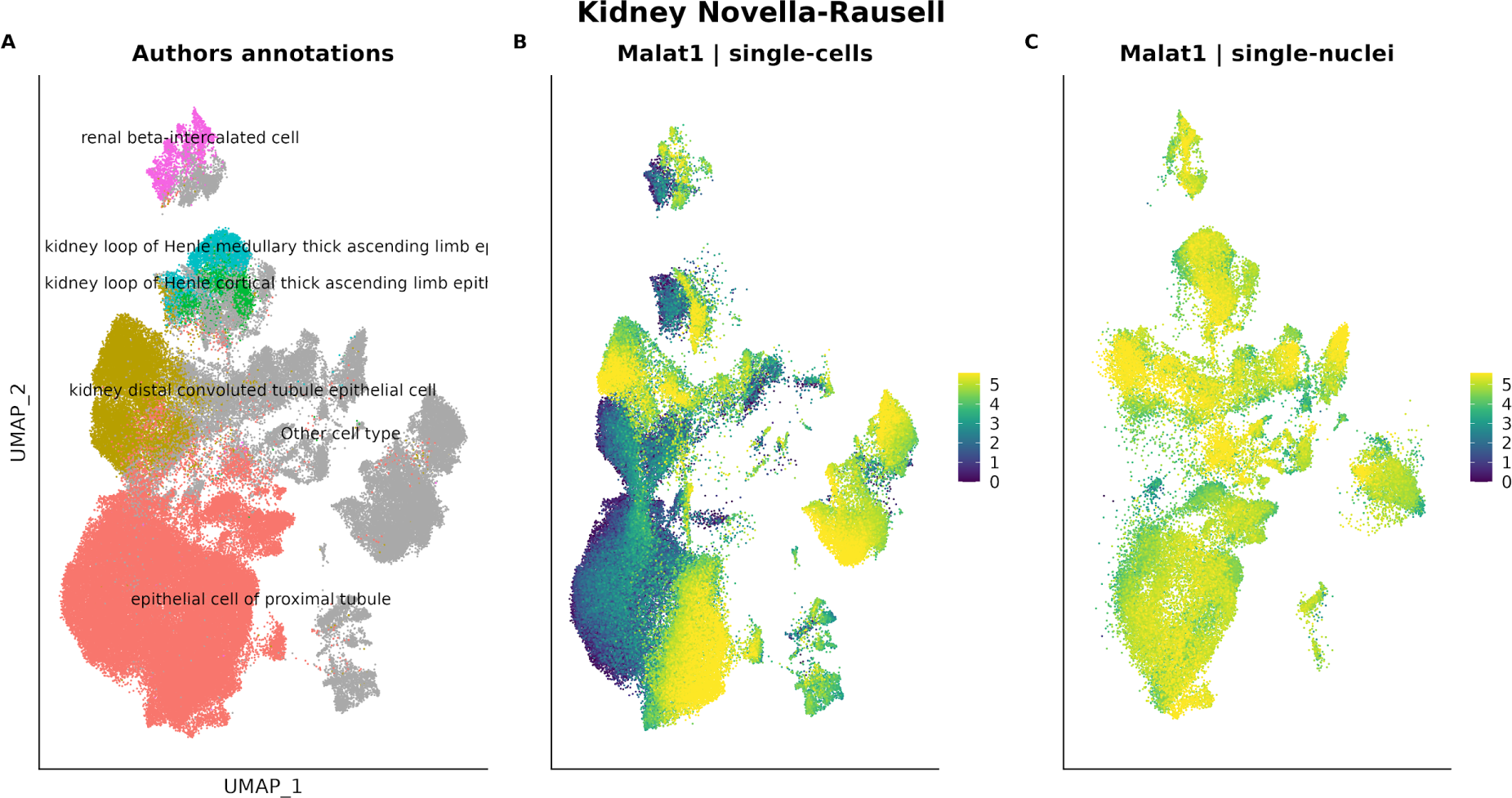
A) Annotated cell types and Malat1 expression of the Novella-Rausell et al. (2023) kidney dataset (single-cells and single-nuclei). Cell types having Malat1+ and Malat-clusters are in colour, the rest of cell types (in grey) have been grouped as “Other cell type”. B) Malat1 expression in the single-cell subset. C) Malat1 expression in single-nucleus subset.

Of note, the distribution of MALAT1 expression in the single-cell subset showed a multimodal distribution. On the other hand, the single-nucleus subset showed a unimodal distribution (Figure 4A). Given that the distribution of the single-nucleus subset overlaps one of the peaks of the single-cell distribution, we hypothesized that, in the single-cell subset, the amount of MALAT1 expression reflected nuclei enriched fractions (MALAT1 high), cells (MALAT1 intermediate) and cytosolic debris (MALAT1 null or low). To further support this fact, we studied the mitochondrial percentage distribution of each sub-distribution found in the single-cell subset. As expected, the distribution of the mitochondrial percentage in the single-cell nuclei enriched fraction was very similar to the single-nucleus subset, while the single-cell cytosolic debris and cell fractions showed higher amounts of mitochondrial reads (Figure 4B).

**Figure 4:**
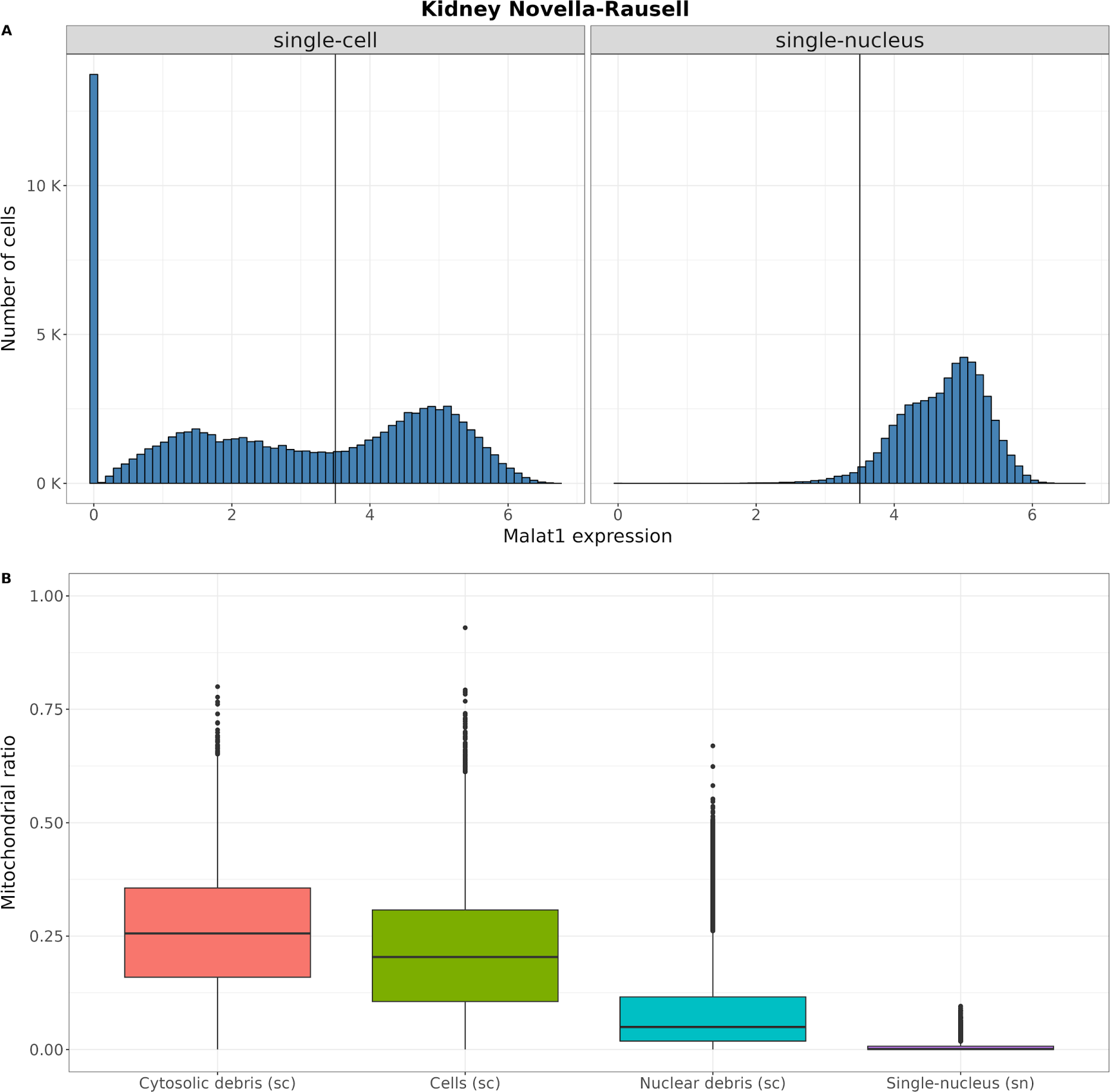
MALAT1 expression distribution in the single-cell and single-nucleus subsets and its relation with the percentage of mitochondrial mRNAs in the Novella-Rausell kidney cell atlas dataset. A) Distribution of MALAT1 expression in the single-cell (left) and single-nucleus (right) subsets. The vertical line indicates the cutoff used to divide the two population of MALAT1+ cells. B) Distribution of the percentage of mitochondrial mRNAs in the three different MALAT1 populations of cells (cytosolic debris, cells and nuclear debris) in the single-cell assay and in the single-nucleus technology.

We then investigated whether the intron fraction of the four single-cell RNA-seq reanalyzed (Tabula Muris kidney 10x and Smart-seq2, MacParland liver and Lukowski retina) allowed to distinguish between the three fractions defined earlier. Two clear populations of barcodes could be easily identified. Those with extremely low introns fraction (the ones that DropletQC characterize as “empty droplet”) and the rest (Figure 5 A-D). The mitochondrial percentage varied among the low introns fraction, compatible with the fact that this subset may represent empty droplets and cytosolic debris droplets that can or cannot have encapsulated mitochondria. On the other hand, the fraction with extremely high intron fraction tended to have less mitochondrial percentage since mitochondrial transcription is not present in the nucleus (same observation for single-cell nuclei enriched subset of the Novella-Rausell kidney cell atlas dataset described above). This is consistent with the classification of these cells done by Braun et al.^21^ that they represent cytoplasm and mitochondrial debris. Table 2 summarizes the characterization of the different cell fractions depending on the relative amount of nUMI, mitochondrial percent and intron content (or MALAT1).

**Figure 5:**
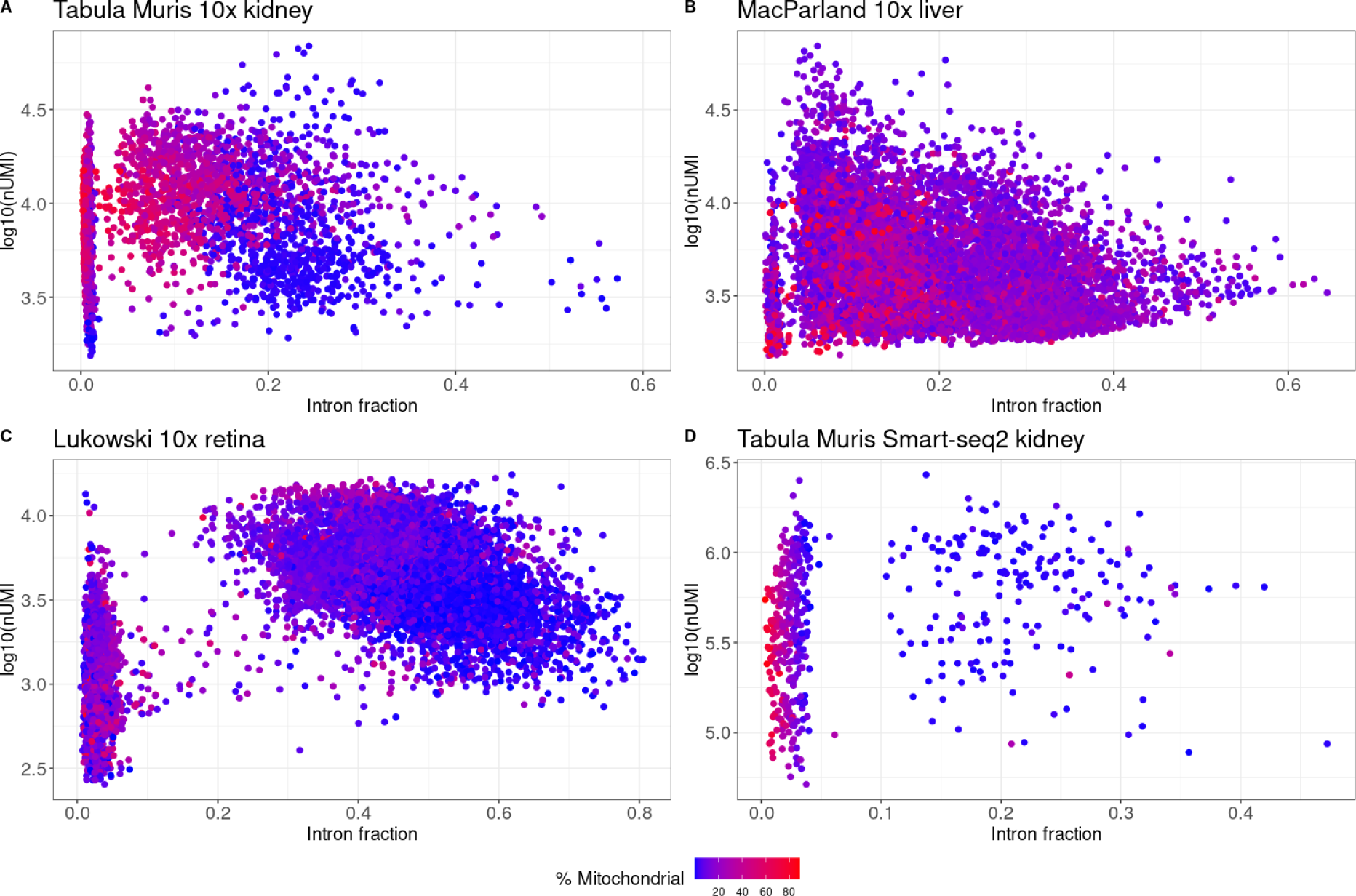
Relationship between intronic fraction (x axis), total UMI (y axis) and mitochondrial content (color scale). A) Kidney 10x Genomics. B) Liver 10x Genomics. C) Retina 10x Genomics. D) Kidney Smart-Seq2.

**Table 2.** Relative amount of intronic content (or MALAT1), mitochondrial and nUMIs of each type of droplet.

| Type of droplet | Intronic/MALAT1 | Mitochondria | nUMIs |
| --- | --- | --- | --- |
| Empty droplet | null | variable | low |
| Mitochondria | null | high | low |
| Cytosolic debris | null/low | high | variable |
| Bare nucleus | high | low | high |
| Damaged/dying cell | high | high | variable |
| Whole cell | intermediate | low/intermediate | high |

### MALAT1-group of cells appears in all tissues from Tabula Sapiens

The Tabula Sapiens dataset allowed us to explore the presence of the MALAT1-group of cells from 28 different tissues and organs from the human body (Figure 6A). In all of them, we identified MALAT1-cell clusters, being kidney, prostate and heart, the three tissues (excluding blood since it contains red blood cells which lack nucleus) with the highest proportion of MALAT1-cells (Table 3 and Supplementary Table 1). Among specific cell types, sperm, kidney epithelial cells, and slow muscle cells have the highest percentage of MALAT1-cells, once erythrocytes were excluded (100 %, 99.412 % and 78.528 %, respectively) (Table 4 and Supplementary Table 1).

**Figure 6:**
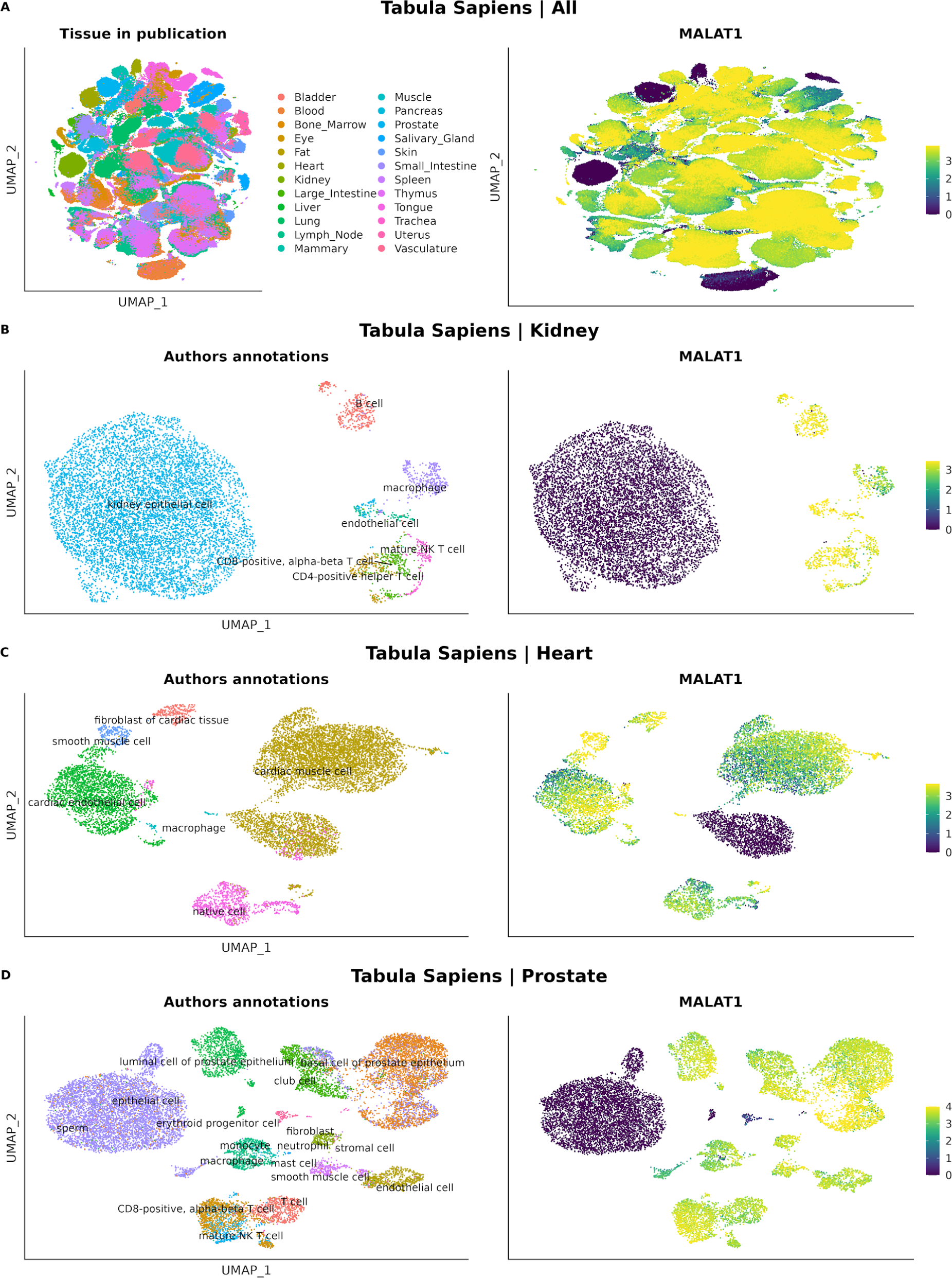
UMAPs showing tissues and their MALAT1 normalized counts in Tabula Sapiens datasets. A) All tissues. B) Kidney. C) Heart. D) Prostate.

**Table 3.** Absolute number of MALAT1-cells and their proportion in the top 5 tissues with more MALAT1-percentage of cells in the Tabula Sapiens dataset.

| Tissue | MALAT1- | MALAT1+ | % MALAT1- |
| --- | --- | --- | --- |
| Kidney | 8301 | 1340 | 86.1 |
| Prostate | 6037 | 10338 | 36.87 |
| Blood | 9824 | 40291 | 19.63 |
| Heart | 1950 | 9555 | 16.95 |
| Tongue | 1990 | 13030 | 13.25 |

**Table 4.** 10 cell types with the highest percentage of MALAT1-cells in the Tabula Sapiens dataset.

| Annotated cell type | MALAT1- | MALAT1+ | Sum | % MALAT1- |
| --- | --- | --- | --- | --- |
| Sperm | 11 | 0 | 11 | 100 |
| Kidney epithelial cell | 8282 | 49 | 8331 | 99.41 |
| Erythrocyte | 9735 | 1269 | 11004 | 88.47 |
| Slow muscle cell | 128 | 35 | 163 | 78.53 |
| Epithelial cell | 7083 | 2079 | 9162 | 77.31 |
| Fast muscle cell | 167 | 185 | 352 | 47.44 |
| Platelet | 103 | 165 | 268 | 38.43 |
| Retinal pigment epithelial cell | 15 | 34 | 49 | 30.61 |
| Cardiac muscle cell | 1782 | 5423 | 7205 | 24.73 |
| Cell of skeletal muscle | 2 | 8 | 10 | 20 |

We further explored the MALAT1 expression of the kidney, prostate and heart datasets from the Tabula Sapiens atlas (Figure 6B-D). MALAT1-cells can be observed as clusters in the UMAPs of the three tissues. In a similar way as in the Novella-Rausell et al. kidney atlas dataset, in the heart dataset the same annotated cell type (cardiac muscle cell) was divided in two clusters, one MALAT1- and another MALAT1+.

On the other hand, the Tabula Muris Senis dataset also revealed the presence of these cells. Among the tissues present in this dataset, pancreas, liver and kidney are those with higher percentage of Malat1-cells (14.6 %, 11.8 % and 6.1 %, respectively) (Supplementary Table 2). The three cell types with the highest proportion of MALAT1-cells are pancreatic acinar cells, ependymal cells and cells of skeletal muscle (51.2 %, 38.2 % and 34.4 %, respectively). Other cell types also affected are those from kidney (collecting duct epithelial cells, 30.2 %, proximal straight tubule epithelial cells, 23.2 %) and heart (cardiac muscle cells, 20.4 %) (Supplementary Table2).

In the same way as we did with the Tabula Sapiens dataset, we explored the UMAPs of the most affected tissues searching for Malat1+ and Malat1-clusters of cells. The original authors divide these datasets into the 10x Genomics and Smart-seq2 subsets. Thus, we explored these subsets separately. In these three tissues (pancreas, liver and kidney), Malat1-cells clustered in the same pattern as for the Tabula Sapiens dataset (Supplementary Fig1 and Supplementary Fig2).

Finally, we confirmed that the presence of MALAT1-cells does not depend on the technology used for sequencing. Both the 10x Genomics (3’ and 5’) and Smart-seq2 subsets showed similar percentages of MALAT1-cells. (Supplementary Table 1 and 2).

## Discussion

Tissue dissociation, a necessary step for acquiring samples suitable for scRNA-seq, subjects cells to traumatic conditions. Previous studies have highlighted the emergence of cell clusters exhibiting up-regulated genes associated with stress conditions^22–25^. However, to date, the presence of cell clusters devoid of immature mRNA content has not been thoroughly investigated as an artifact. While the intronic content metric was initially introduced in 2021 to identify empty droplets in droplet-based scRNA-seq datasets^12^, and later adopted for quality control in heart and brain tissues by two separate publications^21,26^, no studies have demonstrated the widespread presence of cells with null intronic content in publicly available resources such as the human and mouse reference cell atlases. Surprisingly, these seminal findings have received limited attention within the single-cell community, as indicated by the low citation count of the initial study proposing the use of this metric for quality control.

Our reanalysis of single-cell data downloaded from Human Cell Atlas and Tabula Muris, utilizing the percentage of reads mapped to introns as a quality metric, reveals that organs designated as ‘high-quality’ often harbor a variable number of cells with questionable viability. These cells, unnoticed by standard quality control (QC) metrics, are routinely incorporated into downstream analyses. Alternatively, we propose using MALAT1 expression level, given its strong correlation with intronic content, as a new QC metric useful in scenarios where only the gene count matrix is available. Traditionally, MALAT1 has been associated with an inverse correlation to cell health, with higher expression levels observed in dead or dying cells (some tutorials even recommend excluding MALAT1 from analysis). However, our findings reveal an additional usage: negligible or low MALAT1 expression levels correspond to empty droplets or cytosolic debris. Cells with ineffective nuclear lysis could also match this scenario of low MALAT1 and low intronic content.

Those cells presenting high percentage of intronic reads and MALAT1 likely correspond to nuclear debris since their content coincided with the snRNA-seq tissue-matched experiment. On the contrary, nuclei-free cytosolic fractions present null or low levels of intronic reads (or MALAT1) with representative expression of mitochondrial genes. Intact cells, however, show intermediate levels of intronic reads and high UMI counts. This work confirms the presence of artifacts resulting in different cell debris fractions (cytosolic, mitochondrial, nuclear) as described by Braun et al.^21^ in their dataset.

In the kidney and liver datasets, the cell types that showed clusters with no immature mRNA have been already described as difficult-to-dissociate cells. In the kidney dataset, the cells from the proximal tubule are the ones with a higher proportion of cells without immature mRNA, a fact that aligns with what Jansen et al.^27^ recently reported studying the effect of ambient RNA: cells from the proximal tubule are the most vulnerable to the tissue dissociation process. The authors of the liver dataset acknowledged that the cluster of cells labelled as Hepatocyte_6, the one with no unspliced mRNA, had the lowest amount of captured reads and that further investigations were needed to clarify the origin and role of these cells. This, combined with the recent work by Jin-Mi Oh et al.^28^, where they found that hepatocytes are the liver cell type most affected by dissociation effects, seems to indicate that the reduced unspliced mRNA content, not only in this cluster but in all hepatocytes in this dataset, likely comes from cytosolic debris and is a result of the dissociation process.

Although in the publications associated with the kidney and liver datasets, the lack of MALAT1 expression in those cell clusters was not reported, the authors of the retina dataset study did report it and validated experimentally. They hypothesized that the appearance of rods with low MALAT1 expression (which are the same as those with no immature RNA) was due to their degeneration, as the proportion of those cells increased with the post-mortem time of the donor. However, we found that this was not an exclusive feature of rods, as we also found apparent degenerated cones, astrocytes, and microglial cells.

These facts align with the findings of our analysis of the Tabula Sapiens and Tabula Muris Senis datasets which revealed variations in the percentage of MALAT1-cells among different cell types within the same tissue. This variability suggests that the experimental methodologies employed for cell isolation and lysis might not uniformly impact all cell types.

The inadvertent inclusion of low-quality cells in the analysis can yield significant consequences for the outcomes. On the one hand, it may erroneously suggest the presence of specific cell subtypes within a sample, exemplified by the misidentification of the Hepatocyte_6 cluster in the liver dataset. As we noted in this work, usually the same cell type is found in two different clusters, one with introns and the other without introns. This, in turn, may hamper the precise automatic annotation of cell types, and potentially impacts differential expression analysis within clusters of the same cell type or across different cell types. Of particular significance, methods reliant on the relative abundance of spliced (no-introns) versus unspliced (with introns) reads to infer developmental trajectories, such as RNA velocity^29,30^, may yield unreliable results if failing to account for the biases outlined in this study Our work aims to emphasize the importance of the unspliced mRNA fraction metric in the initial quality control of scRNA-seq datasets. As stated before, the use of this metric allows the identification of cells that are likely degraded due to experimental dissociation processes. However, although we showed that it is independent of the sequencing technology, we did not study the effect of different dissociation techniques on the generation of these cells. For example, Denisenko et al. (2020)^23^ reported that the expression of stress-associated genes is much lower if dissociation is carried out in cold conditions. Denisenko et al. (2020) and Jin-Mi Oh et al. (2022)^23,28^ also reported that scRNA-seq and snRNA-seq techniques allow for the recovery of different proportions of each cell type from a tissue, something that agrees with our findings on MALAT1 expression pattern in the kidney dataset from Novella-Rausell et al. (2023). Other studies also evaluated the use of transcription inhibitors^31^ or, more recently, advanced dissociated tissue preservation^32^. Thus far, the presence of stressed cells has been recognized as a significant artefact across various publicly available datasets. However, the quality assessment of these works did not inspect their datasets for the presence of cells with null intronic content.

In summary, our findings reveal that clusters of cells lacking immature mRNA or MALAT1 represent a prevalent artefact in both droplet-based and plate-based scRNA-seq datasets, underscoring the importance of addressing this issue to avoid bias in downstream analyses. Despite the rapid advancements in scRNA-seq technology, the recommended quality control measures have remained largely unchanged since the inception of the technique. With growing evidence suggesting the inadequacy of conventional quality control standards for certain tissue samples, the inclusion of metrics such as the unspliced mRNA fraction becomes imperative.

### Future perspectives

Reference cell atlases, representing both human and mouse body organs, stand as invaluable resources for the single-cell community. However, they often contain numerous low-quality cells that have eluded detection. The presence of artefacts in reference single-cell datasets dampens their quality, potentially compromising the performance of emerging generative single-cell foundational models. These models, pretrained on a vast amount of data (e.g., scGPT^33^ used more than 30 millions of cells from CELLxGENE) rely on the availability of high-quality data for accurate predictions and interpretations. While laboratory protocols strive to yield only high-quality cells, it is imperative that robust bioinformatics strategies are implemented to effectively flag and exclude low-quality cells from analyses. In this way, advancements in single-cell spatial technologies hold promise for mitigating cell loss and enabling the acquisition of whole transcriptome imaging at true single-cell resolution. Adopting these developments will be pivotal in advancing on the reliability of the single-cell genomics field.

## Data availability

All single-cell and single-nuclei RNA-seq datasets used in this study are publicly available from different sources as described in *Materials and Methods*.

## Code availability

The R script to generate the figures of the analysis can be found at https://github.com/funcgen/single_cell_atlases_quality_assessment.

## Author contributions

T.M-A and A.E-C conceived the project. T.M-A performed the bioinformatic analyses. T.M-A and A.E-C wrote the manuscript. A.E-C supervised the project.

## Supplementary Figures legends

Supplementary Figure 1: UMAPs showing tissues and their MALAT1 normalized counts in Tabula Muris Senis datasets. A) All tissues. B) Pancreas (10x Genomics). C) Kidney (10x Genomics). D) Liver (10x Genomics).

Supplementary Figure 2: UMAPs showing tissues and their MALAT1 normalized counts in Tabula Muris Senis datasets. A) All tissues. B) Pancreas (Smart-seq2). C) Kidney (Smart-seq2). D) Liver (Smart-seq2).

## Supplementary Tables legends

Supplementary Table 1: Absolute number and proportion of MALAT1- and MALAT1+ cells in each tissue and cell type contained in the Tabula Sapiens dataset. In the whole dataset and split by sequencing technology (10x 3’, 10x 5’ and Smart-seq2).

Supplementary Table 2: Absolute number and proportion of MALAT1- and MALAT1+ cells in each tissue and cell type contained in the Tabula Muris Senis dataset. In the whole dataset and split by sequencing technology (10x 3’ and Smart-seq2).

