## Supplementary Figures for "Revealing the Prevalence of Suboptimal Cells and Organs in Reference Cell Atlases: An Imperative for Enhanced Quality Control"

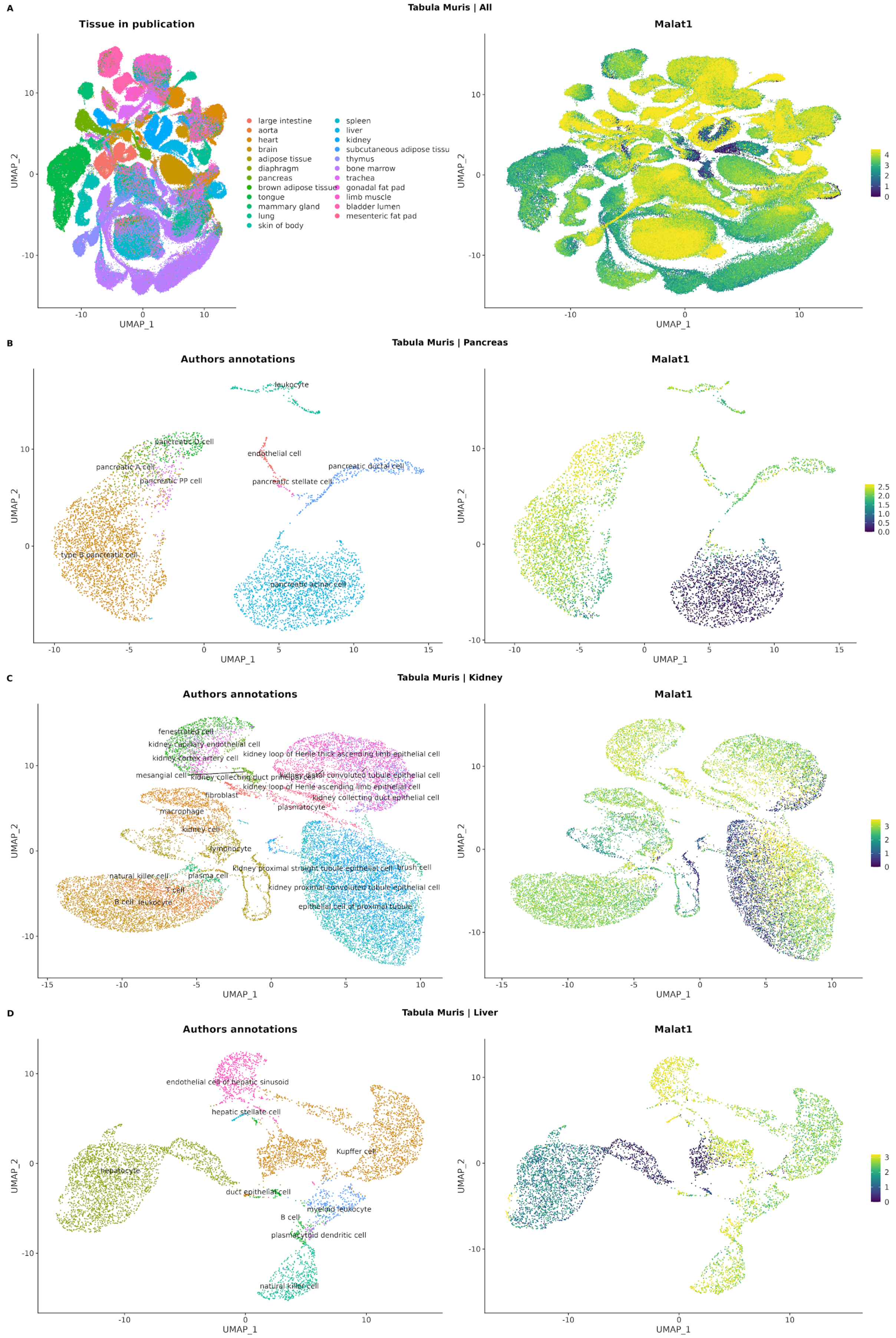


Supplementary Figure 1: UMAPs showing tissues and their MALAT1 normalized counts in Tabula Muris Senis datasets. A) All tissues. B) Pancreas (10x Genomics). C) Kidney (10x Genomics). D) Liver (10x Genomics).

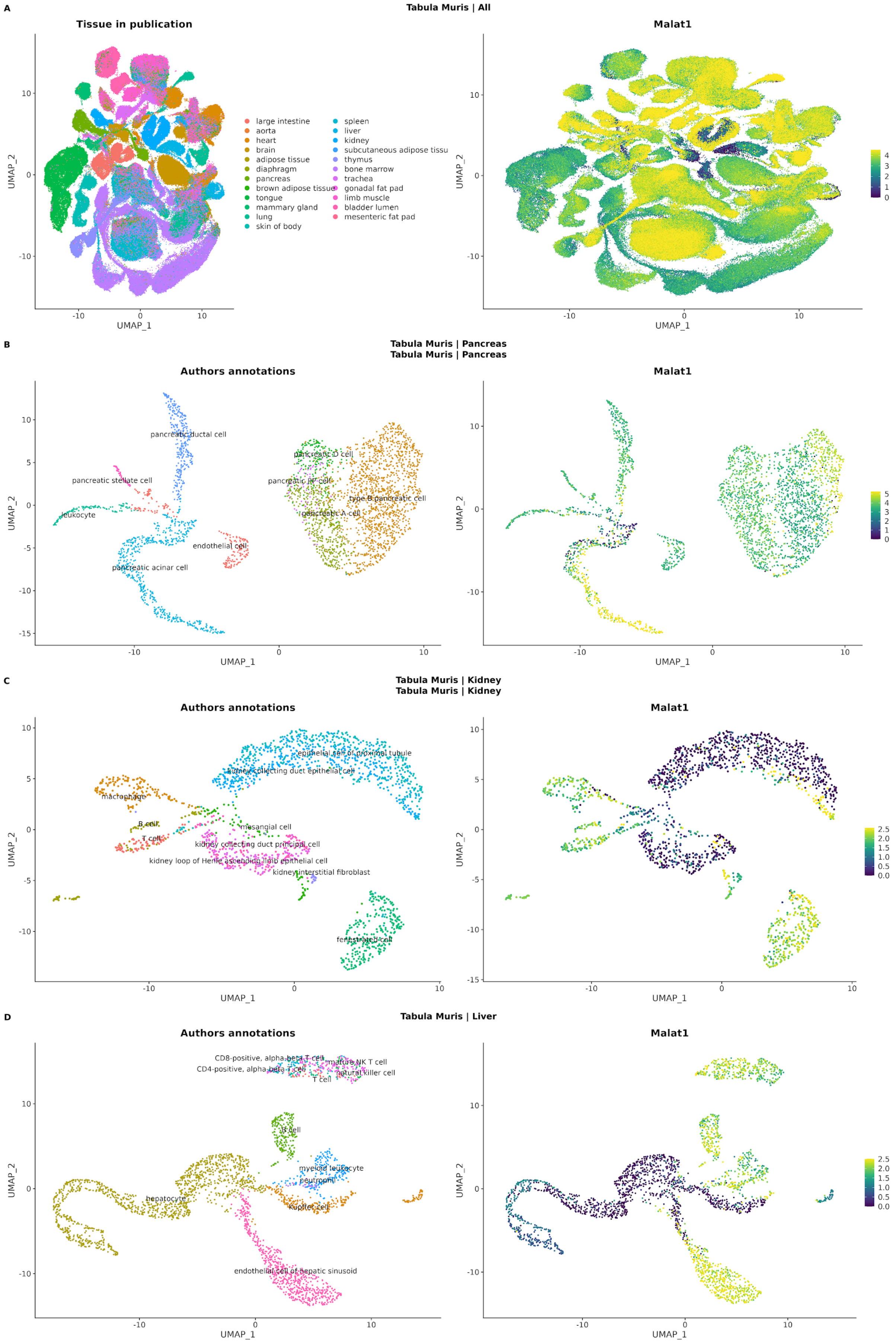


Supplementary Figure 2: UMAPs showing tissues and their MALAT1 normalized counts in Tabula Muris Senis datasets. A) All tissues. B) Pancreas (Smart-seq2). C) Kidney (Smart-seq2). D) Liver (Smart-seq2).
